# Metabolic overlap in environmentally diverse microbial communities

**DOI:** 10.1101/653881

**Authors:** Eric R. Hester, Mike S.M. Jetten, Cornelia U. Welte, Sebastian Lücker

## Abstract

The majority of microbial communities consist of hundreds to thousands of species, creating a massive network of organisms competing for available resources within an ecosystem. In natural microbial communities it appears that many microbial species have highly redundant metabolisms and seemingly are capable of utilizing the same substrates. This is paradoxical, as theory indicates that species requiring a common resource should outcompete one another. To better understand why microbial species can co-exist, we developed Metabolic Overlap (MO) as a new metric to survey the functional redundancy of microbial communities at the genome scale across a wide variety of ecosystems. Using metagenome-assembled genomes, we surveyed over 1200 studies across ten ecosystem types. We found the highest MO in extreme (i.e., low pH/high temperature) and aquatic environments, while the lowest MO was observed in communities associated with animal hosts, or the built/engineered environment. In addition, different metabolism subcategories were explored for their degree of metabolic overlap. For instance, overlap in nitrogen metabolism was among the lowest in Animal and Engineered ecosystems, while the most was in species from the Built environment. Together, we present a metric that utilizes whole genome information to explore overlapping niches of microbes. This provides a detailed picture of potential metabolic competition and cooperation between species present in an ecosystem, indicates the main substrate types sustaining the community and serves as a valuable tool to generate hypotheses for future research.

## Introduction

Microorganisms drive global biogeochemical cycles, but they do not work or live in isolation. In order for any living species to survive they must engage in competition for space and resources with other organisms that share similar nutritional requirements. The concept of loss of species less adapted relative to their competitors is known as competitive exclusion (Gause 1934). When one species cannot sufficiently persist in a habitat, they become locally extinct. Through selection of traits that reduce the dependence on a common resource, populations may shift towards coexistence. This is known as niche partitioning, whereby competition is avoided through the utilization of different resources (Schoener 1974). Evidence that these ecological and evolutionary forces shape microbial communities is prevalent in literature; however, the strength of these forces varies with the availability of resources (reviewed in (Nemergut et al. 2013).

Describing a niche of an organism has remained challenging ever since the concept first emerged (Hutchinson 1957). Typically, closely related species are thought to share similar niches, assuming their evolutionary relatedness is reflected in their nutritional requirements. Recently, neutral genetic markers have emerged as a proxy to measure species’ divergence on an evolutionary timescale; however, these phylogenetic markers (i.e., 16S rRNA genes) are unsuitable to evaluate differences in the biochemical capacity of the organisms. Whole genomes contain information relevant to the metabolic capacity of a species, which is essential to describe the putative niches a microbial species may occupy. If one were to ask about the overlap of two microorganisms’ niches, it is conceivable that this is akin to asking how similar the two are on a genomic level.

With the continued advancement in high-throughput DNA sequencing, large amounts of genomic data are frequently released and available for public use. Several recent publications have reported thousands of novel bacterial and archaeal metagenome-assembled genomes (MAGs; Anantharaman et al. 2016; Delmont et al. 2018; Parks et al. 2017; Tully, Graham, and Heidelberg 2018). The sequencing data originated from hundreds of studies investigating different ecosystems, such that these genomes represent a diverse set of taxa from ecosystems around the globe. This presents an opportunity to address the following important questions: how variable is niche overlap in microbial communities across different ecosystems and does the nature of the overlap (i.e., abundance of genes involved in nitrogen cycling) change based on habitat?

In the current study, we surveyed niche overlap in microbial communities by searching for shared pathways in the metabolic reaction network of species within these communities, which we refer to as ‘metabolic overlap’ (MO). This approach was used to investigate two main questions. First, does the degree of niche overlap in microbial communities vary between ecosystems (i.e., do some communities have more species that utilize the same substrates)? Second, how do these microbial communities vary in the degree of overlap of different metabolic categories (i.e., nitrogen or sulfur metabolism)?

We observed patterns of overlap in microbial community members’ metabolism across different ecosystems, which were largely consistent with literature reports. For instance, a low degree of MO was found in microorganisms involved in highly specialized animal host-microbe associations, while aquatic microbes displayed a cosmopolitan repertoire of strategies for nutrient acquisition. These variations seem to be driven by different categories of metabolism, depending on the ecosystem. In addition, we addressed the question of how much the phylogenetic relationship of microbes corresponds to their metabolic overlap. We found that phylogenetic distance between microorganisms was indeed a good predictor for the degree of MO. The strength of this relationship, however, varied between different ecosystems. Generally, survey-based metrics like MO enable observations of global trends and prompt fundamental questions about the biology and ecology of microorganisms.

## Results

### Definition of metabolic overlap

We defined metabolic overlap (MO) as the number of compounds (i.e., reactants) that can be utilized by two organisms based on their shared metabolic network (Figure 1). For example, an organism (Org_1_) that can perform all steps of denitrification from nitrate (NO_3_^−^) to nitrogen gas (N_2_, four reactions in total) shares two reactants with a partially denitrifying organism (Org_2_) that only reduces NO_2_^−^ to N_2_O. This then results in a MO = 2 (ignoring the rest of their metabolism). Conceivably, identifying MO allows a broad identification of species with overlapping niches by counting the compounds that link complimentary metabolic pathways. As the metabolic routes used to degrade certain substrates can vary between organisms, counting the number of shared reactants will reveal MOs that would not be uncovered by shared reactions only. Furthermore, as the number of reactants can vary between reactions, this approach is more sensitive in identifying weak metabolic similarities between organisms.

**Figure 1.**
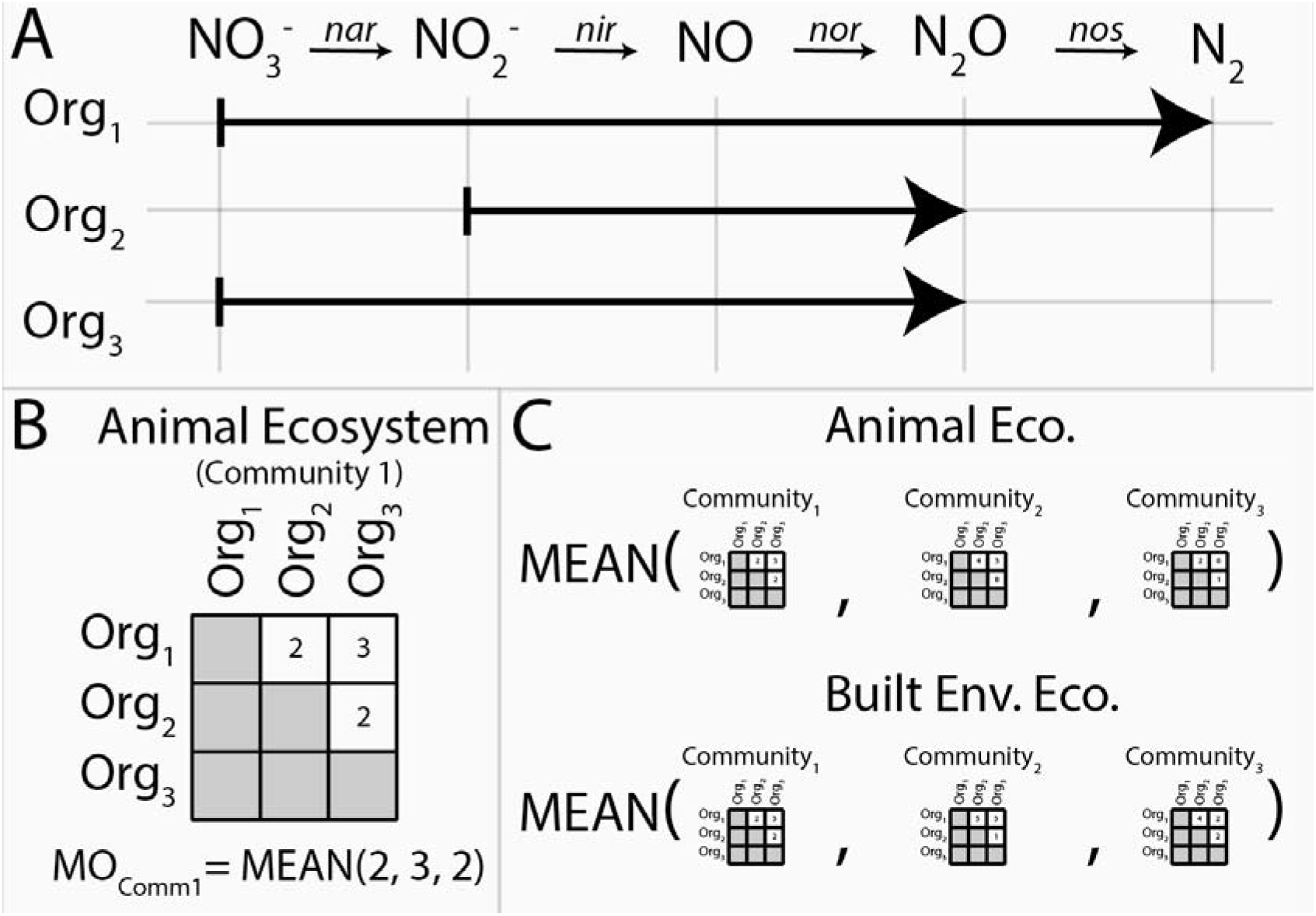
Metabolic overlap is a metric that compares the overlap in the metabolism of two organisms by calculating the number of reactants these species can utilize in common. This is determined by establishing their shared biochemical pathways (A). The number of substrates shared between a set of organisms is represented in a matrix (B), typically a symmetrical distance matrix. The average metabolic overlap of all communities from a given ecosystem are calculated and can be then compared to other ecosystems as seen in the current study (C).

We acknowledge that previous efforts to predict microbe-microbe interactions within microbial communities have been made with similar logic to the current approach. In particular, the NetCooperate software, utilizing the NetSeed framework, is a method to identify putative interactions in a community. It does so by using genome information to predict auxotrophies of the organisms present, based on the incompleteness of certain biosynthesis pathways leading to a dependency of the respective organism to external sources of the lacking metabolite (Levy *et al*., 2015; Carr and Borenstein, 2012). Thus, the NetSeed/NetCooperate approach predicts complementarity between species, which consequently occupy distinct niches, while the goal of our MO approach is to identify to what extent two species fill a common niche.

### Metabolic overlap of microbial communities in different ecosystems

In order to survey the degree of MO in various ecosystems from around the globe, thereby identifying the degree in which microbial species within the community overlap in the niches they fill, the set of Uncultivated Bacteria and Archaea (UBA) MAGs published by Parks and colleagues was utilized (Parks et al., 2017). The average predicted genome completeness of these MAGs ranged from 50-100%. A completion-based inclusion threshold of MAGs was found to have a negligible impact on the average MO of communities (Supplemental Figure 1). In contrast, the number of MAGs included drastically decreased as a result of a more stringent threshold on genome completeness, resulting in ecosystems poorly or not at all represented (Supplemental Figure 1). Thus, we included all 7903 MAGs from the Parks et al. dataset, representing 1248 studies. Studies were classified into their respective ecosystems of origin based on information included in the submission to the public repository or by manual curation if this information was insufficient. This resulted in ten ecosystem categories, with studies that could not be reasonably identified classified as “Other” (Table 1).

**Table 1.** Number of studies and metagenomes within each ecosystem.

|  | Number of studies | Number of metagenomes |
| --- | --- | --- |
| Animal | 130 | 1823 |
| Brackish | 17 | 66 |
| Built | 446 | 1275 |
| Engineered | 122 | 1374 |
| Extreme | 44 | 156 |
| Fresh Water | 59 | 231 |
| Marine | 311 | 1811 |
| Other | 35 | 928 |
| Plant | 3 | 16 |
| Soil | 81 | 223 |
| Total | 1248 | 7903 |

In a given ecosystem, metabolic overlap and the predicted average genome sizes of MAGs were strongly correlated (Supplemental Figure 2; p < 0.01). In addition, average genome sizes significantly varied between ecosystems (Supplemental Figure 3; ANOVA; F = 88; p < 0.001). The average predicted genome sizes were the highest in studies from the built environment (4Mbp +/− 0.65Mbp) and lowest in extreme environments (2Mbp +/− 0.96Mbp; Table 2). The number of MAGs in a given community (grouped per study) negatively correlated with the average MO of the community (Figure 2; Kendall’s tau = −0.38; p < 0.001). As we were interested in investigating how MO varied between ecosystems, irrespective of the differences in genome sizes between ecosystems, we normalized MO to the average genome size of the respective study. Furthermore, the values were scaled so that the average MO of all ecosystems combined was 0 (Figure 3).

**Table 2.**
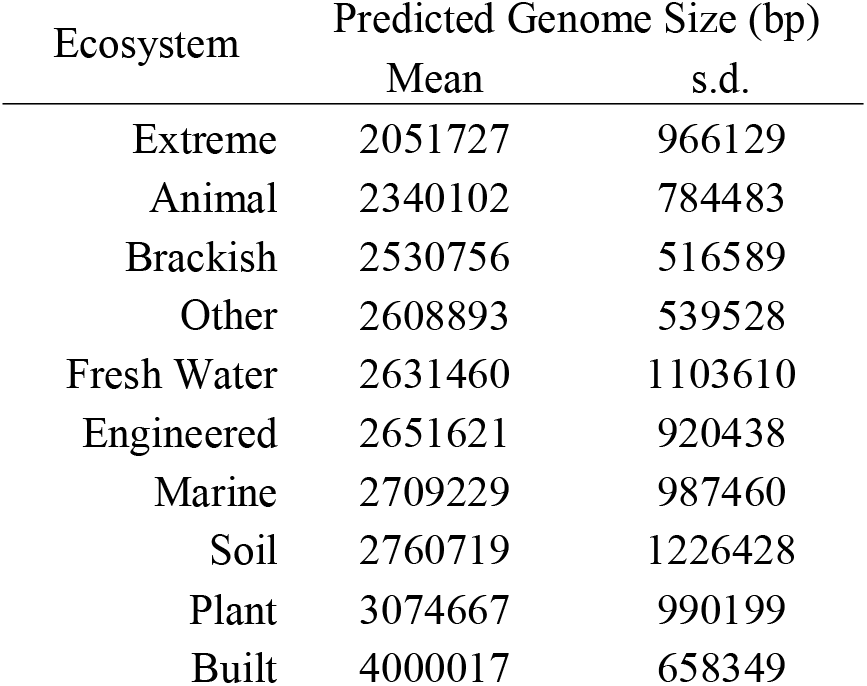
Mean genome size in each ecosystem.

**Figure 2.**
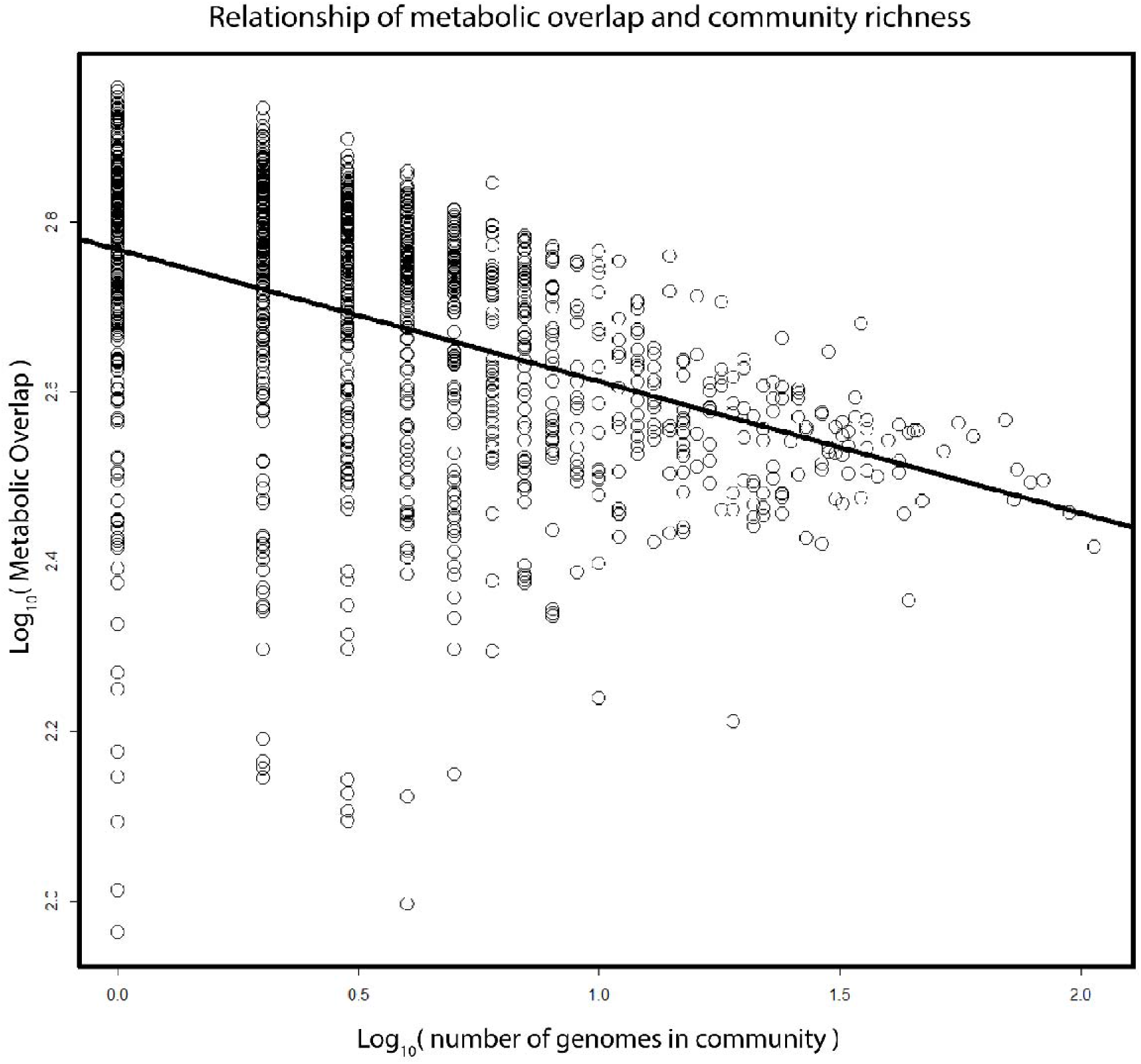
Relationship between metabolic overlap and the number of genomes in a community. Each circle represents one of the 1248 studies. The x-axis depicts the total number of MAGs in a given study, the y-axis the mean metabolic overlap of that study.

**Figure 3.**
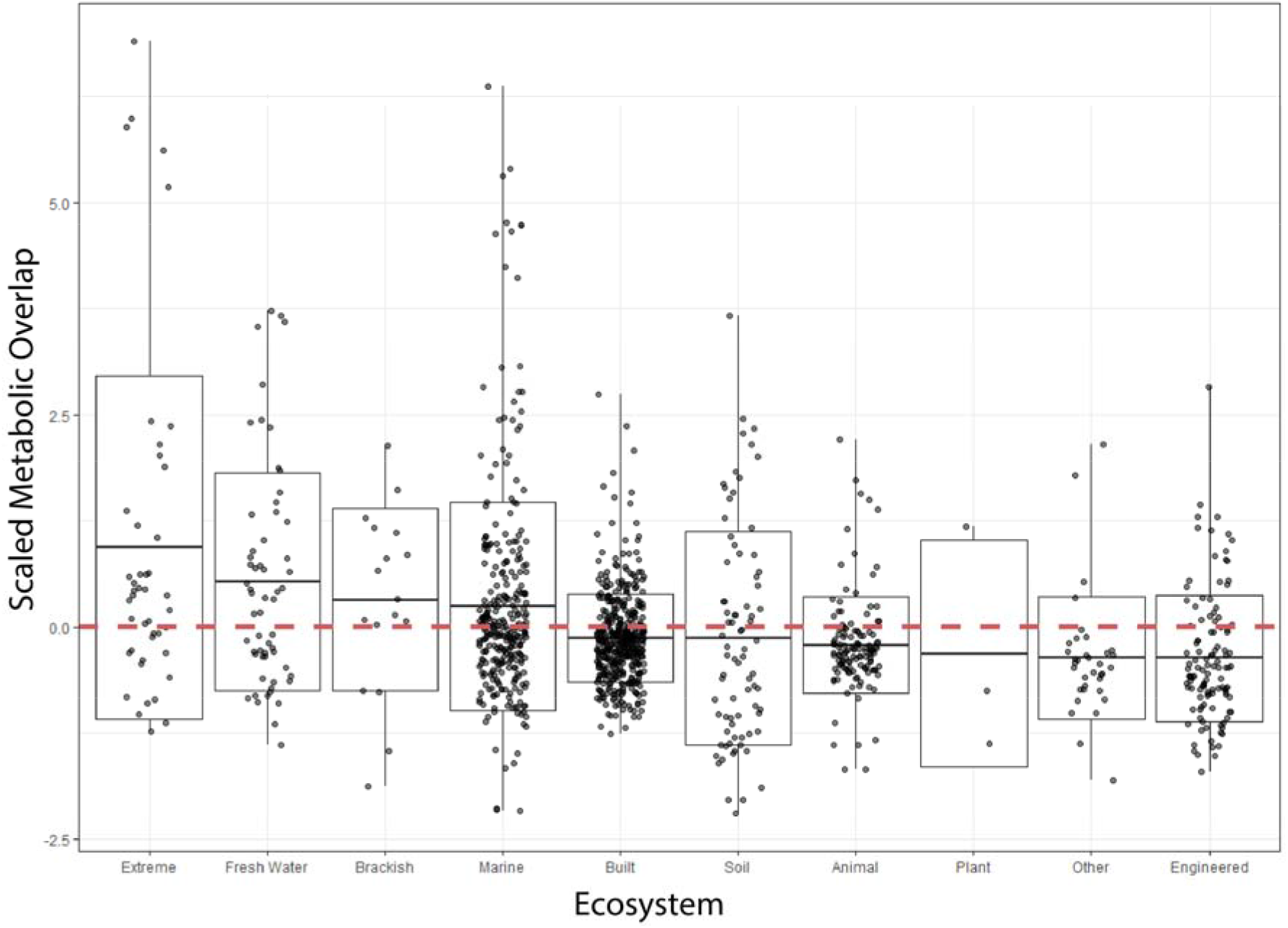
Metabolic overlap across all ecosystems. Boxplots are plotted with the black bar representing the mean, the box is the 25% and 75% quartiles, and the whiskers are the extreme values. A horizontal red dashed line was plotted to indicate 0, which corresponds to the average MO of all ecosystems combined. Each point represents the mean metabolic overlap of all MAGs from a given study.

To evaluate how the MO of microbial communities varied between ecosystems, we determined how the average MO of a single ecosystem differed from the average MO of all ecosystems. Communities from Animal, Built, and Engineered ecosystems had significantly lower MO than average (t-test; p < 0.01; Table 3; Figure 3). On the contrary, those from Extreme, Freshwater and Marine ecosystems had significantly higher MO than average (t-test; p < 0.01; Table 3; Figure 3).

**Table 3.**
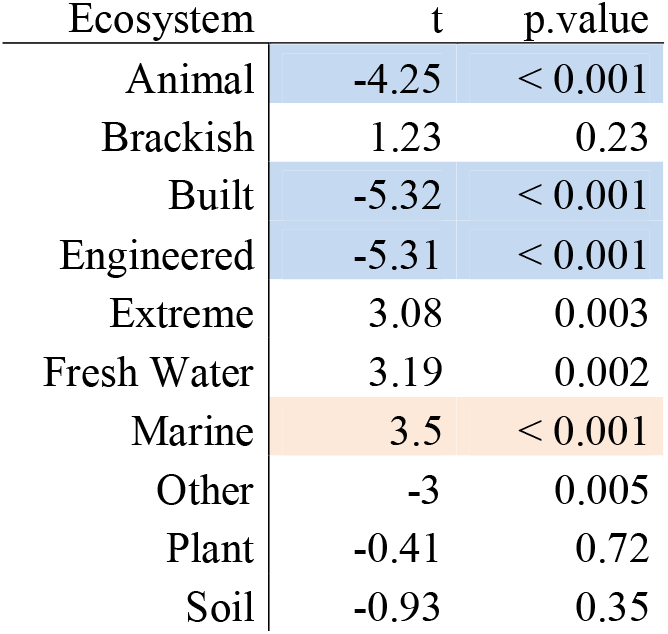
Metabolic overlap statistics in each ecosystem.

### Breakdown of MO scores across different ecosystems to different levels of metabolism

To investigate how metabolic overlap varied between ecosystems within different categories of metabolism (SEED subsystems), the MO within these subcategories was determined for each ecosystem and compared to the average value of all ecosystems (Table 4). Animal, Built and Engineered ecosystems were generally below the average MO for the majority of subcategories of metabolism with a few exceptions (t-test; p < 0.01; Table 4). Communities from Engineered ecosystems had an above average MO in Protein and Nucleotide sugar metabolism, as did communities from Animal ecosystems. In addition, communities from the Animal ecosystem had an above average MO in Nucleotide metabolism. While most subcategories of metabolism from the Built environment were below the average MO, these communities contained higher MO in Nitrogen and Sulfur metabolism (Table 4). In contrast to the above communities, which were dominated by lower than average MO scores, Extreme, Freshwater, and Marine ecosystems had higher than average MO scores in the majority of the categories of metabolism (Table 4).

**Table 4.** Metabolic overlap in different categories of metabolism

| Statistic (t) | Animal | Brackish | Built | Engineered | Extreme | Fresh Water | Marine | Other | Plant | Soil |
| --- | --- | --- | --- | --- | --- | --- | --- | --- | --- | --- |
| Amino Acid Metabolism | -18.808 | 1.0396 | -7.0897 | -8.0706 | 4.3911 | 3.0921 | 7.8829 | -2.8291 | -0.1535 | 0.0306 |
| Aromatic Metabolism | -14.002 | -0.2595 | -4.3238 | -2.9174 | -1.4082 | 1.0256 | 7.0192 | -0.793 | -2.968 | 1.4364 |
| Carbohydrate Metabolism | -10.015 | 0.5472 | -4.4021 | -7.0678 | 3.4987 | 1.9514 | 6.0391 | -4.0708 | -0.6076 | -0.1959 |
| Cofactor Metabolism | -23.996 | 1.7968 | -8.3088 | -9.3914 | 3.4899 | 3.6088 | 8.5258 | -3.1922 | -0.1503 | 0.5693 |
| Fatty Acid Metabolism | -6.2286 | 3.0776 | -7.0429 | -5.054 | 2.5558 | 2.4734 | 6.5712 | -1.5586 | -0.5669 | -2.5669 |
| Nitrogen Metabolism | -15.495 | 1.7972 | 6.9147 | -4.0639 | 2.3216 | 2.2063 | -0.9558 | -0.4684 | -0.4381 | -1.4858 |
| Nucleotide Metabolism | 4.6226 | -0.4585 | -16.574 | 0.0291 | 3.241 | 1.4079 | 4.8181 | -1.164 | -0.6831 | -0.8228 |
| Nucleotide Sugar Metabolism | 3.073 | 1.4103 | -12.295 | 2.9834 | 3.4546 | 3.6569 | -1.2075 | 0.6512 | -0.0622 | 2.3715 |
| Phosphorous Metabolism | -3.8878 | 0.8176 | -8.3411 | -1.0578 | 0.7587 | 2.2691 | 4.6001 | -1.064 | -0.5048 | -0.8658 |
| Protein Metabolism | 3.7253 | 1.8878 | -33.882 | 2.2077 | 5.5824 | 3.9273 | 4.4379 | -0.6715 | 0.2008 | 2.5578 |
| Respiration | -15.021 | 3.4966 | -3.8864 | -5.1202 | 2.8844 | 2.5968 | 9.519 | -1.5019 | -1.0597 | -1.8178 |
| Sulfur Metabolism | -24.636 | -0.3785 | 14.3902 | -10.8816 | -0.6835 | 0.5794 | 1.7196 | -7.2878 | -1.7582 | -2.4058 |
| Secondary Metabolism | -0.41 | 1.005 | -15.753 | -1.161 | 3.738 | 2.241 | 4.644 | 0.609 | NA | 2.243 |

The nitrogen metabolism was used to further investigate the influence of incomplete pathways on the MO. Therefore, the ratios of complete to incomplete denitrifiers were calculated for all ecosystems (i.e., complete denitrifiers encoding all proteins required for NO_3_^−^, NO_2_^−^, NO, and N_2_O reduction; incomplete denitrifiers missing at least one gene; Figure 4A). The Built environment showed the largest MO in nitrogen metabolism and also had the highest ratio of complete to incomplete denitrifiers compared to all other ecosystems (Figure 4B). Contrary, the Animal ecosystem, which by far had the lowest MO in this category also contained mostly incomplete denitrifiers.

**Figure 4.**
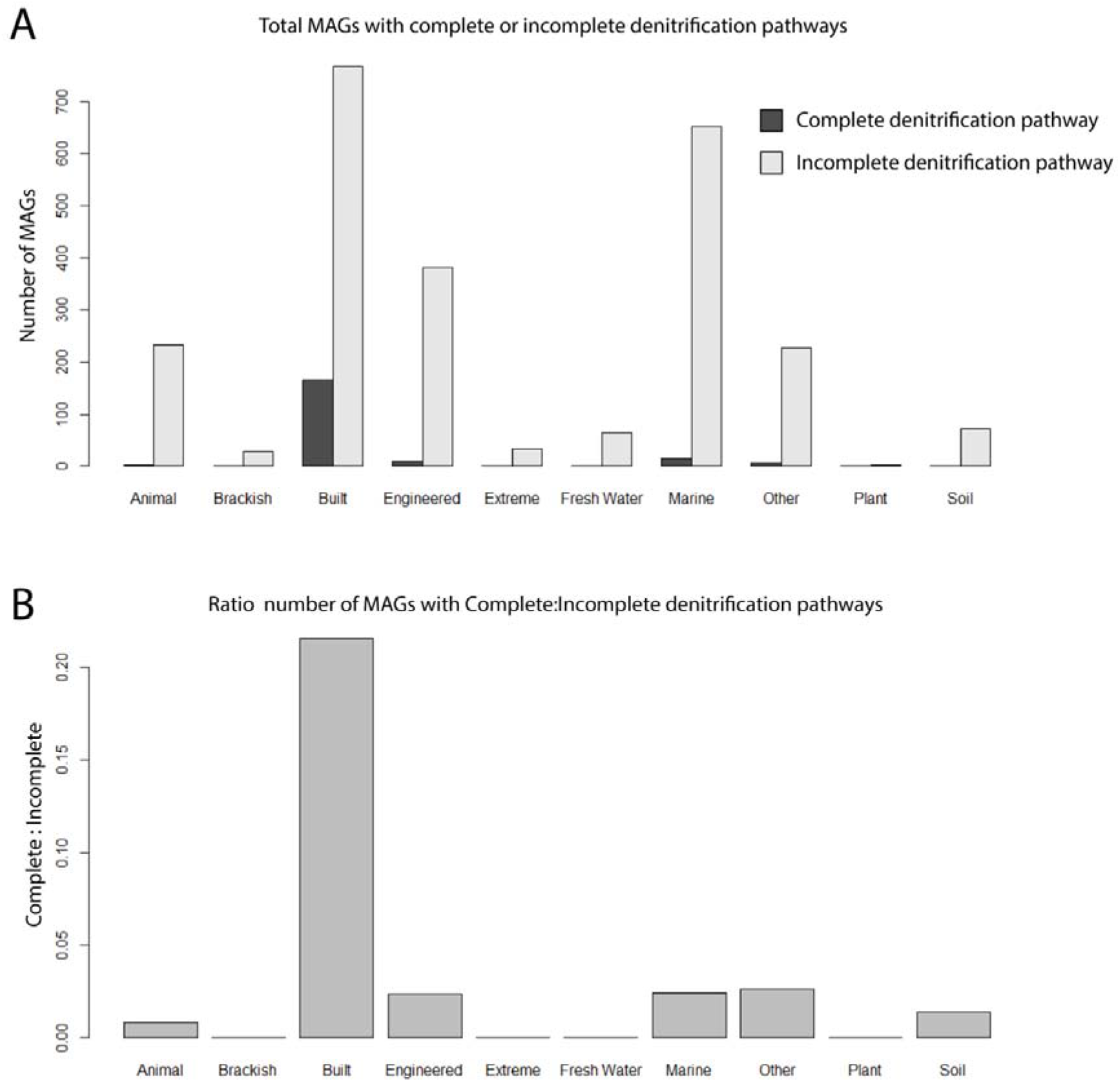
Proportion of complete to incomplete denitrification pathways across different ecosystems. (A) Number of MAGs encoding all proteins to reduce NO_3_^−^ to N_2_ (complete denitrifiers) compared to the number of MAGs with one or more of the respective genes missing. (B) Ratio of complete to incomplete denitrification pathways.

### Phylogenetic relationship of organisms and its relationship to the metabolic overlap

In order to determine if the evolutionary relatedness between MAGs was correlated with MO, the UBCG pipeline was utilized to infer a phylogenetic tree based on a concatenated alignment of 92 universal bacterial marker genes (Na et al. 2018). A significant negative correlation was observed between phylogenetic distance and metabolic overlap for all ecosystems (Figure 5; r =−0.33; p < 0.001), however the strength of this association varied. Phylogenetic distance and MO had the strongest association in Plant (r = −0.64), Built (r = −0.53) and Marine ecosystems (r = −0.47), whereas the lowest associations were seen in Animal (r = −0.16), Extreme (r = −0.19) and Fresh Water ecosystems (r = −0.21; Figure 6).

**Figure 5.**
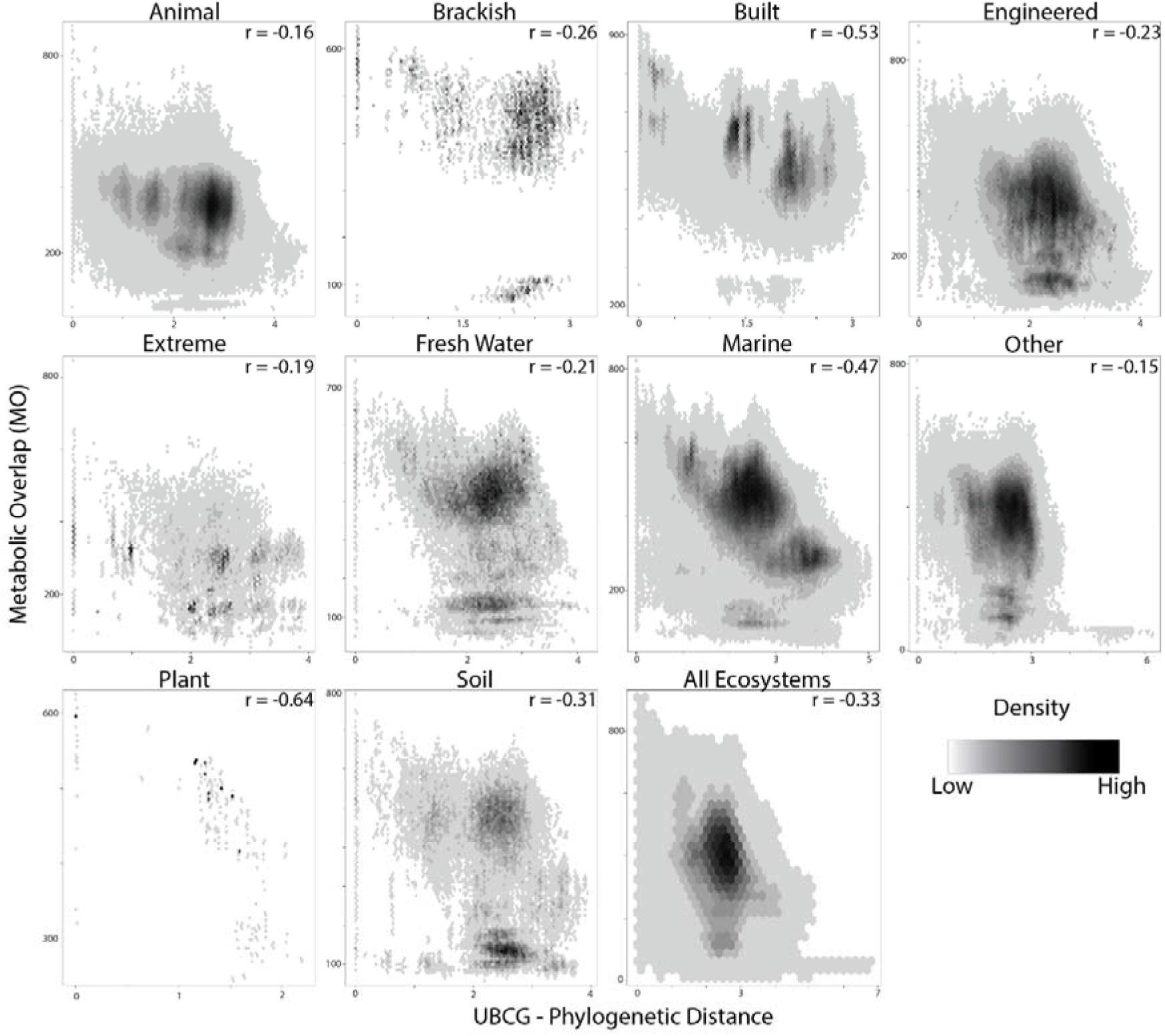
Relationship between metabolic overlap and phylogenetic distance of MAGs. Each point represents a pairwise comparison between two MAGs. The density of points is represented by a black and white gradient. The Spearman’s correlation coefficient is indicated in the upper left-hand corner of each plot.

## Discussion

In the current study a new metric termed MO, which describes how similar two species’ metabolisms are, was developed in the context of a genome-based survey of microbial communities from diverse ecosystems. High MO between two species suggests that they have the capacity to perform similar metabolic reactions, thus have similar growth requirements and fill similar niches. In contrast, low MO suggests that the two species in question may compete for fewer resources. We determined the average metabolic overlap of all community members (i.e., the average MO of all pairwise species comparisons) for a given study, which were grouped into distinct ecosystems based on their origin for comparison (Figure 3; Table 3). The average MO of a community can be similarly interpreted as the pairwise species comparisons. In the case of high average overlap, many community members are overlapping in their biochemistry and could in theory compete for a similar niche, whereas a low average MO would suggest the opposite.

### Ecological and evolutionary drivers of metabolic overlap

There are several well studied ecological forces that shape microbial community structure. Community diversity is maintained via dispersion (immigration and emigration) as well as speciation and extinction. In studying patterns of microbial biogeography, dispersion limitations were seen as one of the driving forces in structuring microbial community patterns in salt marshes and rice paddies, and likely have an influence on the genomic adaptations of marine microorganisms (Kelly et al. 2014; Lüke et al. 2014; Martiny et al. 2006). Microbial biogeography theory has also been applied to help understanding compartmentalized host-associated microbial communities such as microbes in the human lungs (Whiteson et al. 2014). In this study, we observed major ecosystem-dependent differences in the MO of microbial community members (Figure 3; Table 3). This variation may in part be attributed to dispersion limitations inherent to each ecosystem, where ecosystems in which the dispersion of microbial community members is limited would have less overlap than open homogenous ecosystems. Accordingly, the highest MO was observed in aquatic ecosystems, namely communities from the marine open ocean environment, while animal host-associated communities contained some of the lowest MO (Figure 3; Table 3). Ecosystems such as the ocean are likely to not have as strong dispersal limitations as ecosystems like the animal gut or human lungs, and these differences may be a driving force in structuring the MO of their respective microbial communities.

In addition to dispersion as an ecological force, disturbances to ecosystems can also play a large role for species diversity, driving extinction or speciation within the community (Buckling et al. 2000; Connell 1978). Varying degrees of disruption would impart some signature on the metabolic pathways represented in the microbial community. A higher frequency of disturbance would contribute to the extinction of species and reduce the number of redundant metabolisms in a given system. For example, disturbances associated with the marine ecosystem (high MO) such as storms or temperature anomalies are likely less frequent and intense than the regular consumption of foodstuff or intermittent bouts of inflammation in animal guts (low MO) (David et al. 2014; Kashyap et al. 2013; Reese et al. 2018).

### Substrate spectrum as a possible driver of metabolic overlap in ecosystems

The availability of resources, both in quality and quantity, drives which species can thrive in a given system. In the open ocean, the input of labile organic matter is a major factor controlling microbial activity in the photic zone, where phototrophs fix large quantities of inorganic carbon, making new organic matter available to heterotrophic organisms (Aylward et al. 2015; Hansell and Carlson 2002). It is understood that differences in the composition of dissolved organic matter (DOM) enrich for different clades of microorganisms and that the composition of the community is highly influential on the capacity to degrade this carbon (Nelson et al. 2013; Solden et al. 2018). It would follow that a higher substrate selection would drive diversity in the microbial community, and the higher diversity of substrates would then lead to more diverse microbial metabolisms. In the current study, a negative relationship between the richness of a community (number of genomes in a given sample) and their average MO was observed, which suggests that in more diverse communities there is less metabolic overlap (Figure 2). Indeed, there are many studies that report species-specific differences in the composition of host-associated microbial communities ranging from plants to animal hosts (Berg et al. 2014; E.R. Hester et al. 2016; Reese et al. 2018). These differences are in part attributed to the selection of organic compounds that are shared from host to symbiont (Lee et al. 2016; Sasse, Martinoia, and Northen 2018; Zhalnina et al. 2018).

In addition to the quality of substrates, the quantity of organic matter also drastically differs between ecosystems. The concentration of DOC can vary greatly in aquatic systems, with around 40 μmol l^−1^ DOC in groundwater and 5000 μmol l^−1^ in swamps and marshes (Søndergaard and Thomas 2004). Likewise, variations in animal’s diet influence the availability of different substrates for microorganisms. In particular the diet of an animal influences the availability of nitrogen to microbes in animal guts (Reese et al. 2018). Equally, N availability has a strong impact on plant-soil feedbacks, influencing the abundance and metabolism of microorganisms in the rhizosphere (Eric R Hester et al. 2018). If substrates are available in high enough concentrations, the effect of competition may be reduced, potentially leading to a higher number of species consuming a common substrate (i.e., higher MO). In the current study, we observe microbial communities from animal ecosystems had the lowest overlap in categories of metabolism involved in nitrogen and amino acid metabolism, which corresponds to the idea of N limitations in the animal gut and known auxotrophies (Table 4; Reese *et al*., 2018; Zengler and Zaramela, 2018). In contrast, microbial communities from the built environment tend to have higher overlap in nitrogen and sulfur metabolism, though the built environment is a loosely defined ecosystem with limited literature detailing nutrient fluxes through the system (Table 4; Adams et al. 2015). This stark contrast of nitrogen metabolism overlap between the Built and Animal ecosystems, which both generally displayed a lower than average MO, corresponded to the observed number of species capable of complete denitrification. The Built ecosystem had the highest nitrogen metabolism MO, which largely was attributed to the highest proportion of microbial species capable of complete denitrification (Figure 4). This was contrasted by the low number of complete denitrifiers in the animal system. While the differences here could be due to nutrient availability, one should also consider possible differences in life strategies for persisting in a particular environment (i.e., detoxification versus energy conservation).

### Influence of phylogenetic relationship on metabolic overlap

Populations that become isolated and diverge on an evolutionary timescale do so as a result of being exposed to different environments and thus different selection pressures on specific traits, although some mechanisms exist that make this divergence less clear (i.e., convergent evolution, horizontal gene transfer, etc.). In the current study, a relationship was observed between the MO of species and their relatedness (Figure 5), with a reduction of MO with increasing taxonomic distance. While this corresponds well to theory, the strength of the relationship between phylogenetic relatedness and MO varied between ecosystems, suggesting that ecological differences between these ecosystems influence this relationship.

The dominant taxonomic groups often vary between different ecosystems as a result of the underlying nutrient profiles or physical properties of those ecosystems. This may be a result of stronger selection pressures in a given ecosystem for traits specific to a few select monophylogenetic groups (i.e., methanogenesis, ammonia and nitrite oxidation), as opposed to traits that are more widespread (i.e., denitrification). Phylogenetic groups may vary in the number of traits (i.e., some groups are more metabolically versatile than others), and MO is determined by the number of reactions a given pair of species share. For example, Zimmerman et al., found that a set of phylogenetically diverse Bacteria and Archaea had the potential to produce a subset of three extracellular enzymes (Zimmerman, Martiny, and Allison 2013). Specifically, the ability to produce these enzymes was non-randomly distributed phylogenetically. It follows that ecosystems which have strong selection pressures for metabolically diverse phylogenetic groups would have a weaker relationship between the phylogenetic relatedness and metabolic overlap.

### Caveats and limitations of genetic predictions of metabolic overlap

The emergence of vast amounts of sequence data has allowed the assembly of genomes of microorganisms from fragmented DNA isolated from the environment. The degree of information in whole genomes compared to that from marker genes (both phylogenetic and metabolic) is likely to provide significant advances in our understanding of the genetic organization of microorganisms. In addition, knowing that a certain set of genomes were physically in the same sample is advantageous in addressing fundamental questions about the ecology and evolution of microbial communities from natural settings. Unfortunately, there are still significant limitations when dealing with metagenome-assembled genomes. Specifically, the amount of information lost in the process of genome assembly and binning reduces our understanding of population-level genetic variation. Current sequencing depths do not provide sufficient coverage for the metagenomic assembly of low abundance organisms’ genomes, narrowing our view of genetic linkages between species towards the highly abundant species. However, these are mainly technological limitations, with solutions like long read sequencing becoming increasingly more available. Additionally, there is a significant lack of information about the environments in which samples were taken in the public archives, limiting what can be assessed with metrics such as metabolic overlap, and calling for an urgent need to provide as much metadata on samples as possible.

In addition to the technical limitations mentioned above, there are also limitations in methods such as MO, which rely heavily on accurate automated annotation of genetic elements in genomes. Specifically, database quality is a key driver in the accuracy of survey studies such as the one presented here. A major issue is the inability to assign functions to many genes, even in the genomes of the most well studied microorganisms (35% hypothetical proteins in *E. coli* genome; Ghatak et al. 2019). Apart from the limitations to automatic annotation methods, there are different levels of biology associated with niches that are not captured in genome-level information. These limitations include a lack of information of whether a gene is transcribed, whether the transcript is translated to a functional product and ultimately variations in affinity and activity of this protein. The variation in transport efficiency and regulatory mechanisms certainly contributes to the competitive advantage of an organism and thus the niche this organism fills. These complexities are not easily derived from genomic information. Idealistically, as emphasized by (Bowers et al. 2017), in order to improve discovery-based approaches that rely on machine readable formats of public repositories, additional information should accompany MAG submissions. This set of information would not only help assess the quality of the genome but aid in associating the genetic information to the biology and ecology of the organism. Ideally, such information should include conditions of the environment from which the species’ genome was obtained (i.e., pH and temperature), and if the species was cultivated, any physiological parameters that may have been measured (i.e., growth rate, substrate usage profile and affinities, etc.).

## Conclusions

The observation of variation in MO across different ecosystems begs several questions about the nature of microbial community metabolism. Specifically, what drives metabolic versatility in microbial communities? Are there generalizable rules that can be deduced? Survey-based studies enriched with additional information, such as those highlighted above, may shed additional light on important factors that drive MO. In addition, there is a severe need to complement predictions based on the genetics of microorganisms with phenotypic data. Ultimately, understanding drivers of microbial community metabolism will lead to a better ability to predict and engineer microbial communities for industrial or conservational purposes.

## Methods

### Data origin and Annotation of Ecosystems

Metagenome-assembled genomes (MAGs) utilized in the current study comprised the set published by Parks et al. (Parks et al. 2017). The Uncultured Bacterial and Archaeal (UBA) MAGs were downloaded from the author’s repository (https://data.ace.uq.edu.au/public/misc_downloads/uba_genomes/). The accompanying data from the UBA MAG set, including CheckM metrics of predicted genome completeness and size, was obtained from the publication (Parks et al. 2017). Each study in the UBA set of MAGs was manually sorted into a set of nine ecosystems and an unclassifiable category called ‘Other’.

### Metabolic overlap calculation

All MAGs were subsequently annotated using a custom pipeline based on the SEED API (Aziz et al. 2008; Overbeek et al. 2005). In brief, protein encoding genes (pegs) were called from the assemblies using svr_call_pegs (http://servers.nmpdr.org/sapling/server.cgi?pod=ServerScripts). Each of these proteins was then assigned to a figfam with svr_assign_using_figfams. The association of a protein to a biochemical reaction was then made with svr_roles_to_reactions. A custom script (rxn_expandinfo) associated reactions with compounds from the reaction database which is found on the ModelSEED git repository (https://github.com/ModelSEED). Finally, the number of compounds shared between two organism’s set of biochemical reactions is calculated to create a pair-wise MO score, and a distance matrix was constructed to store this information. This was made using the custom python scripts rxn_to_connections and lists_to_matrix, respectively (https://github.com/ericHester/metabolicOverlap). The distance matrix represents the MO of all organisms within a single community and the average MO of all of these organisms is utilized in comparisons in this study.

In addition to an overall MO score for a community, the above approach was used to calculate the MO of various sub-categories of metabolism for the respective community. In addition to the above, an additional step was performed where pegs were assigned to their respective SEED subsystems and filtered with a custom script utilizing svr_roles_to_subsys. With pegs assigned to these metabolic categories, the above pipeline was used to identify reactions and compounds shared between pairs of organisms, subsequently resulting in a distance matrix similar to that above. In this case, the distance matrix stores the MO of the community pertaining to a specific category of metabolism. Matrices and accompanying data were further analyzed in R (R Core Team 2016).

### Relating phylogenetic distances of MAGs to their MO within communities

In order to associate the phylogenetic distance of assembled genomes to their MO, the UBCG pipeline was utilized (Na et al. 2018). This pipeline extracts 92 conserved phylogenetic marker genes and builds multiple alignments for each gene. The resulting alignments are concatenated and a maximum likelihood tree is inferred. This tree was imported into R utilizing the *ape* package and distances were extracted from the tree object with the *cophenetic* function (Paradis, Claude, and Strimmer 2004). The result is a distance matrix containing phylogenetic distances between each pair of MAGs. Subsequently, this phylogenetic distance matrix and the distance matrix storing MO scores were correlated using the *mantel.test* function from the ape package. The Spearman’s rank correlation coefficient was calculated for each ecosystem subset.

## Acknowledgements

We would like to acknowledge Michiel van den Heuvel for contributing to the metabolic overlap pipeline. Funding was provided by the European Research Council (ERC AG ecomom 339880) and the Netherlands Organisation for Scientific Research (NWO; Gravitation Grant SIAM 024.002.002, VIDI grant 016.Vidi.189.050).

---

**Supplemental Figure 1.**
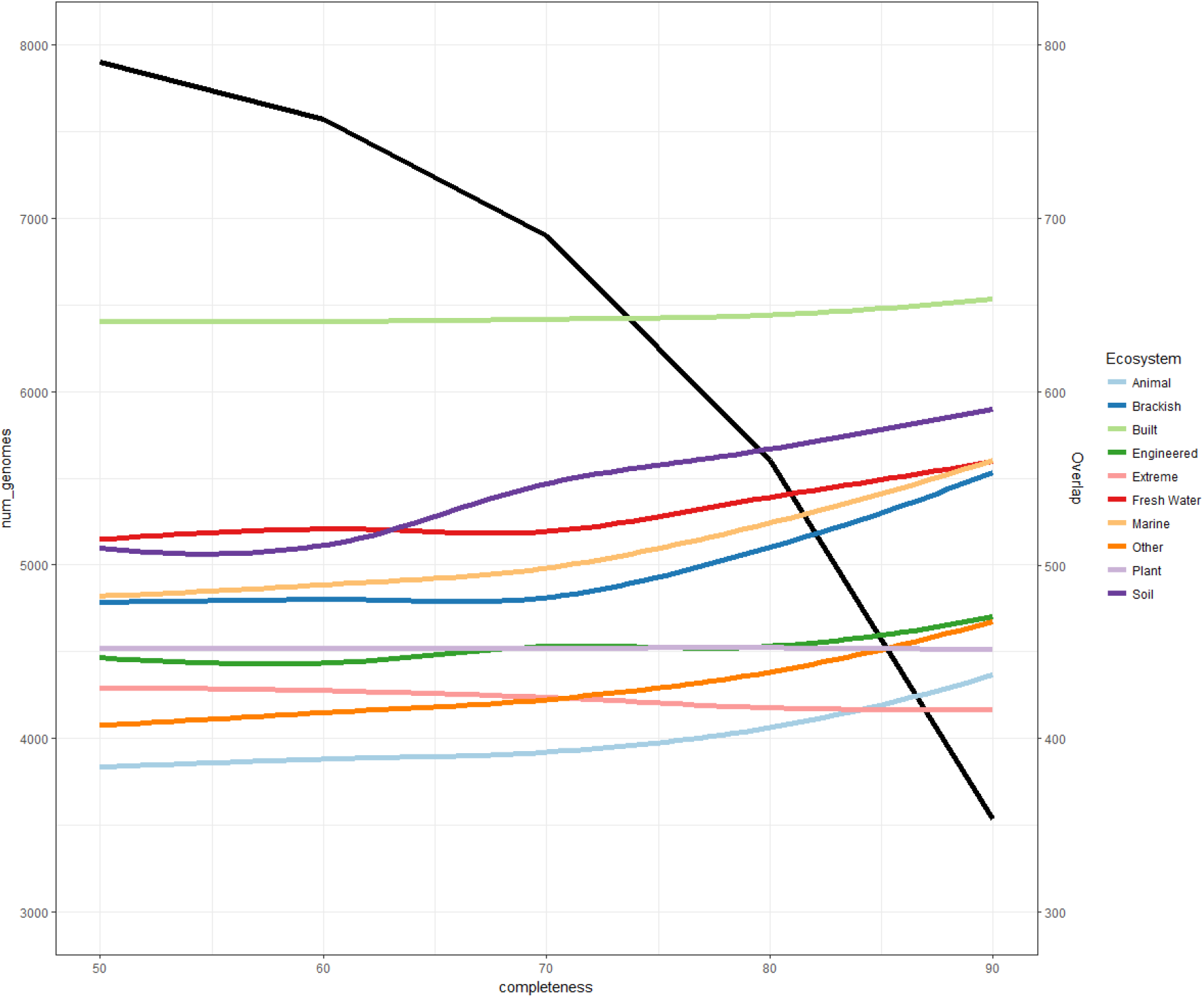
Relationship between the genome completeness and the average metabolic overlap observed (colored lines, right axis). The number of MAGs retained at the different completeness cutoffs is indicated by the black line (left axis).

**Supplemental Figure 2.**
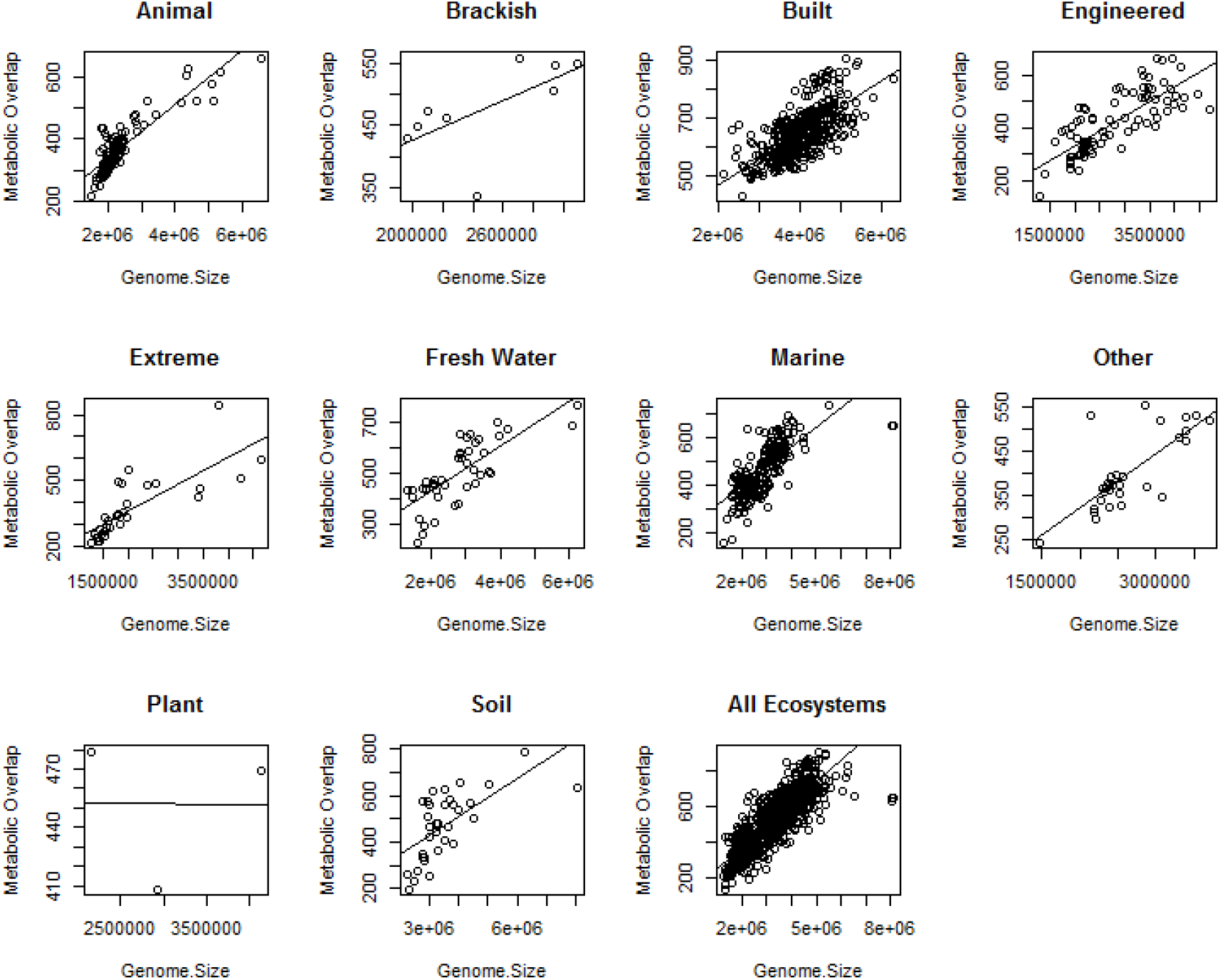
Relationship between metabolic overlap and genome size. Each circle represents one study. The y-axis indicates the average metabolic overlap of all MAGs in one study, and on the x-axis the average genome size for all MAGs in this study.

**Supplemental Figure 3.**
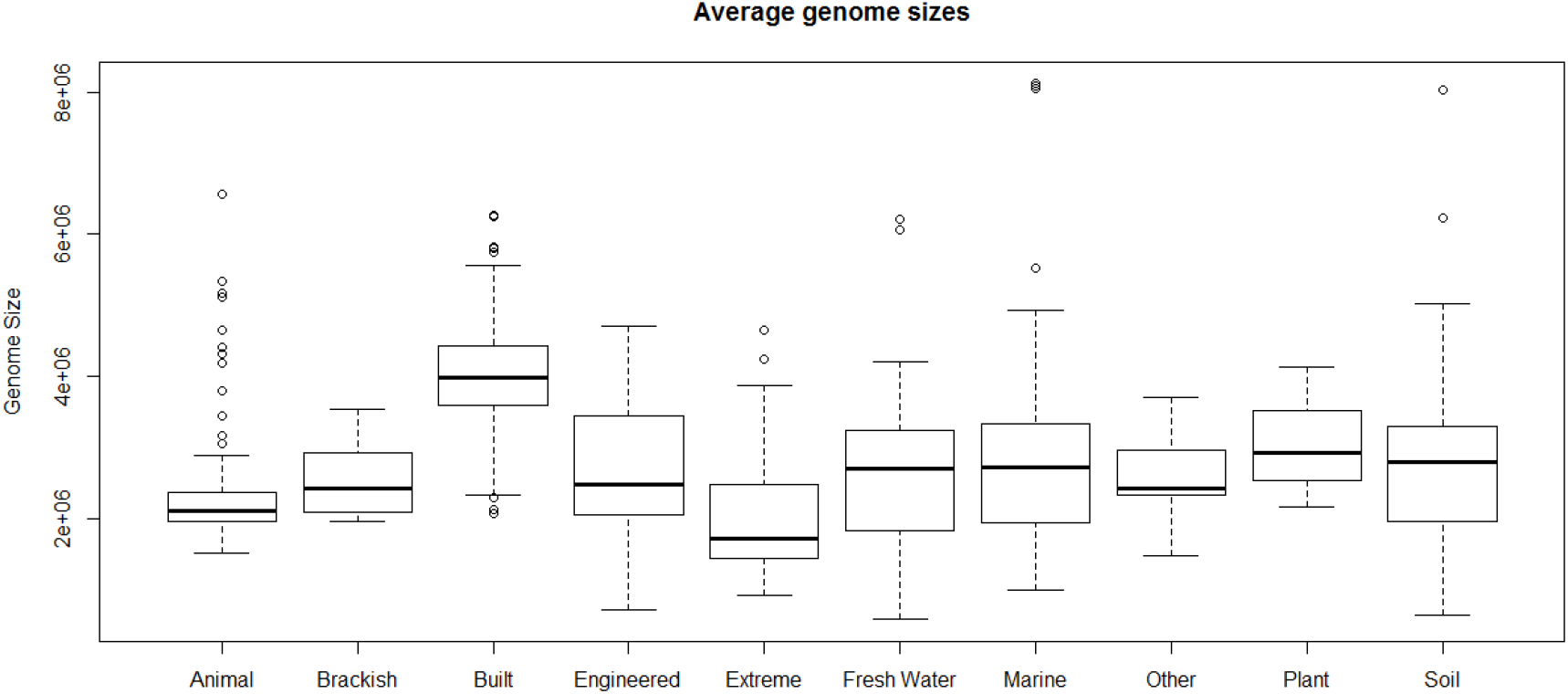
Average genome sizes across ecosystems. The black bar of the boxplot indicates the median, the box edge represents the upper and lower quartiles, whiskers denote extreme values, and individual points are outliers.

